# Opposing signaling pathways regulate morphology in response to temperature in the fungal pathogen *Histoplasma capsulatum*

**DOI:** 10.1101/546853

**Authors:** Lauren Rodriguez, Mark Voorhies, Sarah Gilmore, Sinem Beyhan, Anthony Myint, Anita Sil

## Abstract

Phenotypic switching between two opposing cellular states is a fundamental aspect of biology, and fungi provide facile systems to analyze the interactions between regulons that control this type of switch. A long-standing mystery in fungal pathogens of humans is how thermally dimorphic fungi switch their developmental form in response to temperature. These fungi, including the subject of this study, *Histoplasma capsulatum*, are temperature-responsive organisms that utilize unknown regulatory pathways to couple their cell shape and associated attributes to the temperature of their environment. *H. capsulatum* grows as a multicellular hypha in the soil that switches to a pathogenic yeast form in response to the temperature of a mammalian host. These states can be triggered in the laboratory simply by growing the fungus either at room temperature (where it grows as hyphae) or at 37°C (where it grows as yeast). Prior worked revealed that 15-20% of transcripts are differentially expressed in response to temperature, but it is unclear which transcripts are linked to specific phenotypic changes such as cell morphology or virulence. To elucidate temperature-responsive regulons, we previously identified four transcription factors (Ryp1-4) that are required for yeast-phase growth at 37°C; in each *ryp* mutant, the fungus grows constitutively as hyphae regardless of temperature and the cells fail to express genes that are normally induced in response to growth at 37°C. Here we perform the first genetic screen to identify genes required for hyphal growth of *H. capsulatum* at room temperature and find that disruption of the signaling mucin *MSB2* results in a yeast-locked phenotype. RNAseq experiments reveal that *MSB2* is not required for the majority of gene expression changes that occur when cells are shifted to room temperature. However, a small subset of temperature-responsive genes is dependent on *MSB2* for its expression, thereby implicating these genes in the process of filamentation. Disruption or knockdown of an Msb2-dependent MAP kinase (*HOG2*) and an APSES transcription factor (*STU1*) prevents hyphal growth at room temperature, validating that the Msb2 regulon contains genes that control filamentation. Notably, the Msb2 regulon shows conserved hyphal-specific expression in other dimorphic fungi, suggesting that this work defines a small set of genes that are likely to be conserved regulators and effectors of filamentation in multiple fungi. In contrast, a few yeast-specific transcripts, including virulence factors that are normally expressed only at 37°C, are inappropriately expressed at room temperature in the *msb2* mutant, suggesting that expression of these genes is coupled to growth in the yeast form rather than to temperature. Finally, we find that the yeast-promoting transcription factor Ryp3 associates with the *MSB2* promoter and inhibits *MSB2* transcript expression at 37°C, whereas Msb2 inhibits accumulation of Ryp transcripts and proteins at room temperature. These findings indicate that the Ryp and Msb2 circuits antagonize each other in a temperature-dependent manner, thereby allowing temperature to govern cell shape and gene expression in this ubiquitous fungal pathogen of humans.

## Introduction

The ability of microbes to sense and respond to the environment is key for survival in a variety of niches. Thermally dimorphic fungi make a dichotomous developmental choice depending on their environment. In the soil, they grow in a multicellular, hyphal form (also referred to here as the filamentous form) that also produces vegetative spores. When the soil is disturbed, hyphal fragments and associated spores can be inhaled by mammalian hosts. The body temperature of the host is a sufficient signal to switch the growth pattern of the fungus to a specialized host form (unicellular yeasts in the case of *Histoplasma capsulatum*) that expresses virulence factors associated with survival in the host and pathogenesis. In the laboratory, *H. capsulatum* grows in a hyphal form at room temperature (RT) and switches to a yeast form at 37°C, providing an excellent system to study the regulation of developmental switches that are relevant to disease progression during infection.

*H. capsulatum* is a fungus endemic to the Ohio and Mississippi River valleys where 60-80% of inhabitants test positive for exposure during their lifetime [1]. The transition to the yeast form is thought to facilitate the lifestyle of the fungus in the host, where it replicates within macrophages and causes disseminated disease. Although *H. capsulatum* is ubiquitous, little is known about the molecular regulators and effectors that enable the bidirectional switch between the infectious soil form and the pathogenic host form. A number of studies [2–7] have shown that a significant fraction of the genome is differentially expressed between *H. capsulatum* hyphae grown at RT and *H. capsulatum* yeasts grown at 37°C, including known virulence factors that are specifically expressed at mammalian body temperature. However, little is known about the role of the majority of differentially expressed transcripts in either the biology of the organism or the switch between developmental forms.

In previous work, to elucidate how such a large morphological and transcriptional restructuring occurs in response to temperature, we utilized forward genetic screens to identify four transcription factors, *RYP1-4*, that are required for <u>y</u>east-<u>p</u>hase growth. Mutation of any of the *RYP* genes results in constitutive hyphal growth independent of temperature. Additionally, each of these transcription factors is required for the normal transcriptional response to temperature [2, 7, 8]: the transcriptional profile of the *ryp* mutants at 37°C looks drastically different from wild-type yeast cells but very similar to wild-type hyphae grown at RT [2]. We also found that Ryp transcripts and proteins accumulate to higher levels at 37°C than at RT [2, 4, 7, 8], which presumably contributes to their ability to trigger a temperature-dependent program of gene expression.

Here we elucidate pathways that function at RT to antagonize the Ryp regulatory circuit and prevent yeast-phase growth. We perform a genetic screen for yeast-locked mutants and identify the signaling mucin *MSB2* as required for hyphal formation at RT. Orthologs of Msb2 have been studied in *Saccharomyces cerevisiae* and *Candida albicans* where this transmembrane protein stimulates a number of signaling pathways, including the high osmolarity glycerol (HOG) pathway in response to osmotic stress, as well as the filamentous growth pathway in response to nutrient limitation [9–15]. We examine the role of *MSB2* in *H. capsulatum* as an environmental sensor that controls the morphologic response to RT growth. We find that *MSB2* is required for the expression of a small set of temperature-responsive genes, and that this regulon shows conserved expression in the filamentous form of other dimorphic fungi. To validate that this *MSB2* regulon contains genes that promote hyphal growth, we disrupt both a MAP kinase and APSES transcription factor whose transcriptional induction at RT is dependent on *MSB2* and show that the resultant mutants are defective for normal hyphal growth in response to temperature. Additionally, we provide the first molecular evidence that the Ryp circuit and the Msb2 circuit are mutually antagonistic in a temperature-dependent manner, resulting in the ability of temperature to control cell shape and associated characteristics.

## Results

### The *MSB2* gene is required for hyphal growth at RT in *Histoplasma capsulatum*

Previous studies have identified and characterized genes required for the yeast phase of *H. capsulatum* and other thermally dimorphic fungi [2, 7, 8, 16]. These studies have shed light on the regulation of yeast phase growth; however the regulation of filamentation remains largely unclear. Similarly, how RT growth inhibits the yeast program is also not understood. We sought to shed light on these mechanisms by identifying mutants that inappropriately maintain the yeast state and fail to form hyphae at RT. We used *Agrobacterium*-mediated insertional mutagenesis and screened for mutants that retained their yeast morphology after growth for 6 weeks at RT. One of the mutants we obtained failed to transition to hyphal growth after a shift to RT, which normally induces filamentation (Fig. 1A). Instead, the mutant continued to divide as yeast (Fig. S1). Whole-genome sequencing of the mutant led to the identification of the *Agrobacterium* T-DNA insertion site in the promoter region of the *MSB2* gene (Fig. 1B). Introduction of a wild-type copy of the *MSB2* gene on an episomal plasmid (hereafter referred to as “*msb2* + *MSB2”* or the “complemented strain”) restored the ability of the mutant to form hyphae at RT (Fig. 1C). Knockdown of *MSB2* by RNA interference (RNAi) in a wild-type background recapitulated the yeast-locked phenotype during growth at RT (Fig. 1D). Overexpression of *MSB2* at 37°C resulted in disruption of the normal yeast morphology and the appearance of polarized projections reminiscent of hyphae (Fig. 1E). Taken together, these data indicate that *MSB2* is necessary and sufficient to trigger filamentous growth.

**Fig. 1.**
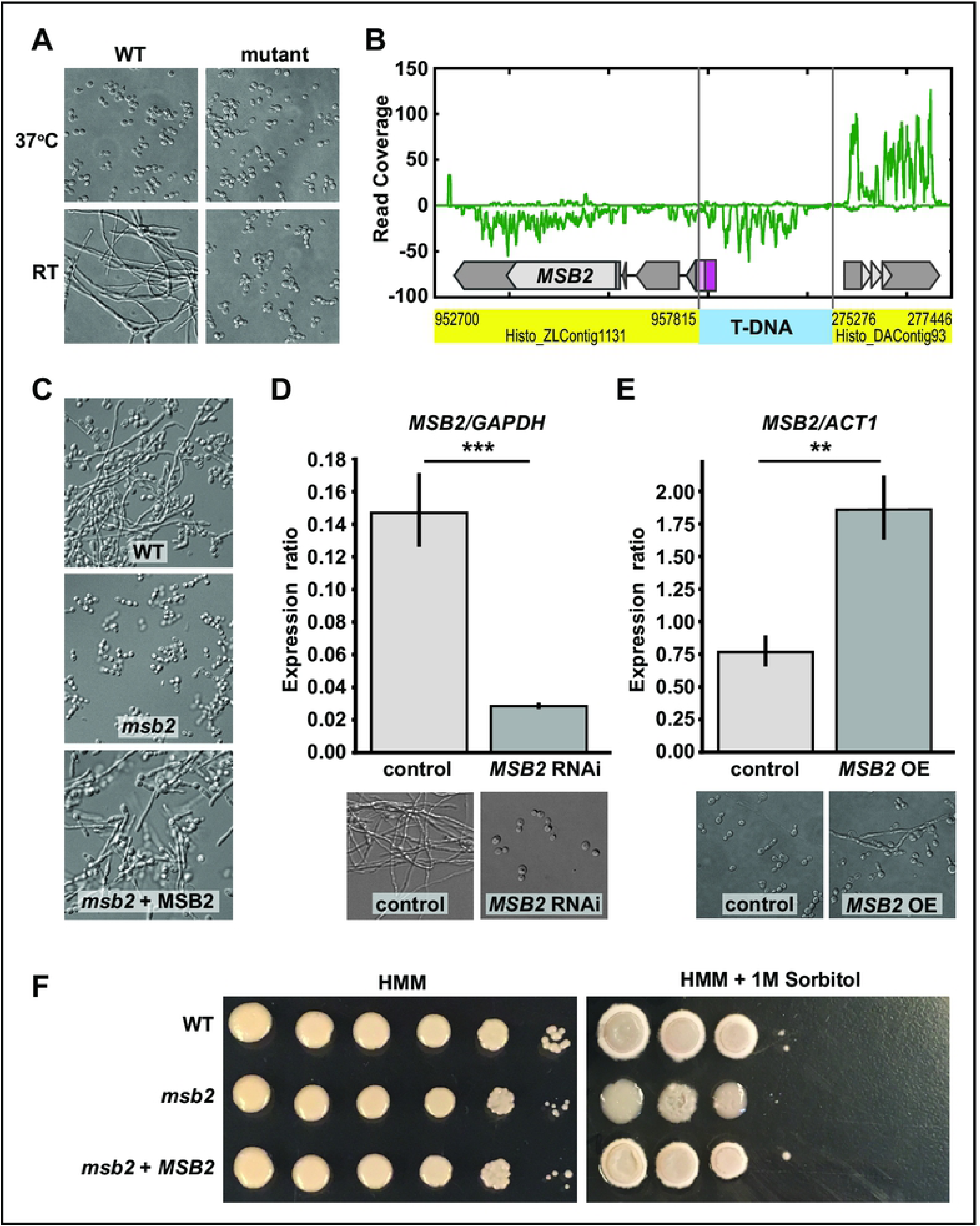
*MSB2* is necessary and sufficient for filamentation of *H. capsulatum* in response to temperature. (A) Cell morphology of wild-type *H. capsulatum* and mutant after 4 days at RT or 37°C in HMM + GlcNAc media. (B) Genome sequencing alignment indicating the transposon insertion site upstream of the *MSB2* gene. This alteration results in transcript initiation in the T-DNA insertion (purple box), thereby introducing an upstream ORF (pink box). Thus the resultant *MSB2* transcript is non-functional because it does not yield a normal translation product. (C) Cell morphology of wild-type *H. capsulatum, msb2*, and *MSB2* complementation strain shown after 8 days at RT. (D) *MSB2* RNAi strain cell morphology is shown after 6 week incubation at RT and the control strain after 4 days at RT. qRT-PCR analysis showed significant depletion of *MSB2* transcript in the RNAi knockdown strain compared to the vector control (P<0.001). (E) *MSB2* overexpression was driven by the *GAPDH* promoter on an ectopic plasmid. qRT-PCR analysis of *MSB2* expression level showed a significant increase in the overexpression (OE) strain compared to control at 37°C (P<0.01). Cells are shown after growth at 37°C for 12 days. (F) Wild-type, *msb2*, and *msb2+MSB2* were grown on HMM solid media +/-1M sorbitol to observe any osmotic stress response that is reflected in the colony morphology at 37°C. HMM colonies were imaged at day 8 after plating and HMM +1M Sorbitol were imaged at day 17.

The function of Msb2 in *H. capsulatum* was previously unknown, though it has orthologs in the distant fungal species *Candida albicans* [10], *and Saccharomyces cerevisiae*, where its function has been more thoroughly examined [9, 11–15]. The *S. cerevisiae MSB2* ortholog acts as a sensor in the osmotic stress response pathway. The response to high osmolarity in *H. capsulatum* has not been thoroughly characterized. To determine if *H. capsulatum MSB2* plays a role in responding to high osmolarity, we plated serial dilutions of wild-type, *msb2*, and the complemented strain on solid HMM media with 1M sorbitol (Fig. 1F). Interestingly, we observed that the wild-type and complemented strains exhibited a wrinkly colony morphology in the presence of sorbitol at 37°C, consistent with some degree of filamentous cells in the colony. In contrast, the *msb2* mutant gave rise to smooth colonies on high sorbitol, correlating with yeast-form cells. These data suggest that the morphology of *H. capsulatum* changes in response to high osmolarity and that this altered cell shape is dependent on Msb2.

### The transcriptional changes elicited by RT growth are largely independent of *MSB2*

Cells lacking *MSB2* are unable to form hyphae at RT. To understand the contribution of *MSB2* to the myriad transcriptional changes that accompany RT growth, we surveyed the transcriptome of wild-type, *msb2*, and complemented strains as they were shifted from 37°C to RT. Yeast cultures were grown in normal yeast-promoting conditions (37°C, 5% CO_2_) and transferred to RT without CO_2_ to promote filamentation. This timecourse was performed in two different carbon sources, glucose (Fig.S2) and N-Acetylglucosamine (GlcNAc) (Fig.2A). We chose GlcNAc because we previously showed that it expedites the transition from yeast to hyphae at RT [17]. Wildtype and the complemented strain produced filaments after 2 days at RT in GlcNAc, but the *msb2* mutant remained in the yeast form over the 8-day timecourse. RNA was isolated from wild-type, *msb2*, and the complemented strains, and we performed RNAseq analysis at day zero (37°C) and 2, 6, and 8 days after the transition to RT. Fig. 2B is a scatter plot showing transcript abundances between two biological replicates of wild-type grown at 37°C, and as expected there is a high Pearson correlation between the samples (r = 0.99) and a corresponding low average deviation from the line of best fit (rmsd = 0.36). In contrast, wild-type hyphae harvested eight days after transfer to RT had an expression profile that was quite distinct from wild-type yeast at 37°C (Fig.2C, r = 0.86, rmsd = 1.08), consistent with the previously observed differences between mature yeast and hyphal cultures [2–7]. Surprisingly, the transcriptional program of the *msb2* mutant, which remained locked in the yeast form even at 8 days post-shift, looked very similar to that of wild-type hyphae (Fig. 2E, r = 0.97, rmsd = 0.57), and very different from wild-type 37°C yeast cultures (Fig. 2D, r = 0.86, rmsd = 1.12) in spite of their common morphology. We conclude from these data that the majority of the temperature-dependent transcriptional program is independent of *MSB2*. Since the *msb2* mutant is yeast-locked, the small set of Msb2-dependent genes may be linked to the ability of *H. capsulatum* to form hyphae.

**Fig. 2.**
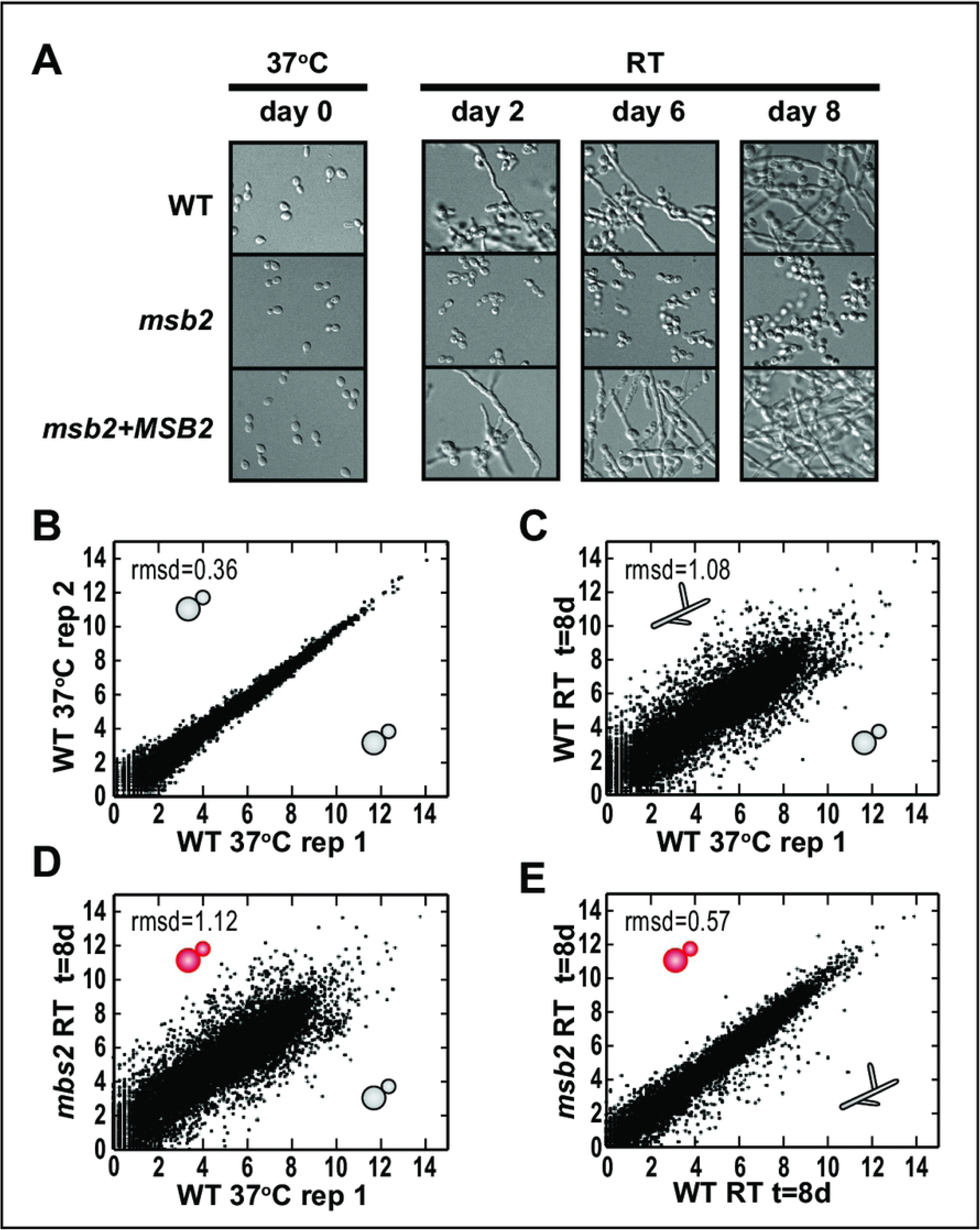
The majority of the temperature-dependent changes in gene expression is independent of Msb2. (A) Experimental setup for RNAseq timecourse. Cells shown were grown at 37°C with 5% CO2 in HMM media with N-acetylglucosamine (GlcNAc) as the carbon source. Once shifted to RT, cells were collected at day 2, 6, and 8 post temperature shift. Images were taken at each time point to compare cell morphology of wild-type, *msb2*, and complemented (*msb2+MSB2*) strains. (B-E) Scatter plots of relative transcript abundances as log2(cpm). (B) Wild-type (WT) 37°C replicate 1 (day 0) vs replicate 2 (day 0). (C) WT 37°C (day 0) vs WT RT (day 8). (D) WT 37°C (day 0) vs *msb2* RT (day 8). (E) WT RT (day 8) vs *msb2* RT (day 8). The morphology of each sample is depicted with a cartoon schematic of yeast or hyphae.

### Transcriptional profiling identified a small set of Msb2-dependent genes with varying functions

As described above, to assess the role of Msb2 in the transcriptional response to temperature, we examined the transcriptional profile of wild-type, mutant, and complemented strains before and after shifting cultures from 37°C to RT in two different media. Liquid cultures were split at t = 0 to give biological replicates, and the entire experiment was carried out on two distinct occasions (batches). Genotype and conditions for each profiled sample are given in S1 Data, with biological replicates collected both at the same time and on different days. There was some variability based on the particular batch, and the transcriptional program in GlcNAc vs. glucose was similar but not identical, with differences that were largely independent of the temperature-dependent changes. To extract the temperature and *MSB2* dependence of each gene from our full data set, we fit a linear model with independent terms for each time point and genotype with additional terms to control for effects due to media (GlcNAc vs. glucose) or batch. Estimated read counts and fit terms are given in S2 Data. We considered genes significantly differential with respect to a fit term if they had at least a two fold change and a p-value less than 0.05 after multiple hypothesis correction. Normalized expression levels of significantly differential genes are given in S3 Data and rendered as a heatmap in Fig. 3B. Consistent with the single sample comparisons (Fig. 2B-E), over a quarter of the transcriptome had significant differential expression over the transition (1870 genes up (11 + 1694 + 165; Fig. 3A) and 1115 genes down (40 + 1067 + 8; Fig. 3A) when comparing the wild-type yeast-phase transcriptome at day 0 (37°C) vs. the wild-type hyphal transcriptome at day 8 (RT)). In contrast, when comparing wild-type day 8 with the *msb2* mutant at day 8, only 3% of the transcriptome was significantly differential (93 genes up in the *msb2* mutant compared to wild-type (11 + 42 + 40; Fig. 3A) and 249 genes down in the *msb2* mutant compared to wild-type (165 + 76 + 8; Fig. 3A), again meaning that the vast majority of transcriptional changes upon shift to room temperature are intact in the *msb2* mutant. For the small subset of genes whose expression is dependent on Msb2, we observed some correlation between morphology and Msb2-dependence, with either (1) increased expression in the yeast-locked *msb2* mutant vs. wild-type hyphae *and* increased expression in wild-type yeast cells vs. wild-type filaments (40 genes; Fig. 3A) or (2) decreased expression in the yeast-locked *msb2* mutant vs. wild-type hyphae *and* decreased expression in wild-type yeast cells vs. wild-type filaments (165 genes; Fig. 3A). This pattern was more common than transcripts that were up in the yeast-locked *msb2* mutant vs. wild-type hyphae *and* up in wild-type hyphae vs. wild-type yeast (11 genes; Fig. 3A) or transcripts that were up in wild-type hyphae vs. wild-type yeast *and* up in yeast-locked *msb2* mutant vs. wild-type hyphae (8 genes; Fig. 3A). This observation is consistent with the hypothesis that genes correlated with (and perhaps required for) hyphal morphology should be upregulated in wild-type hyphae over wild-type yeast as well as upregulated in wild-type hyphae over the yeast-locked *msb2* mutant. Similarly, genes that correlate with yeast morphology should have the opposite expression pattern. Therefore, we further inspected these jointly regulated “filament-associated” (165 genes; Fig. 3A) and “yeast-associated” (40 genes; Fig. 3A) gene sets for potential morphological regulators, noting that a subset of these genes could control morphology and others could be effectors associated with a particular morphologic state.

**Fig. 3.**
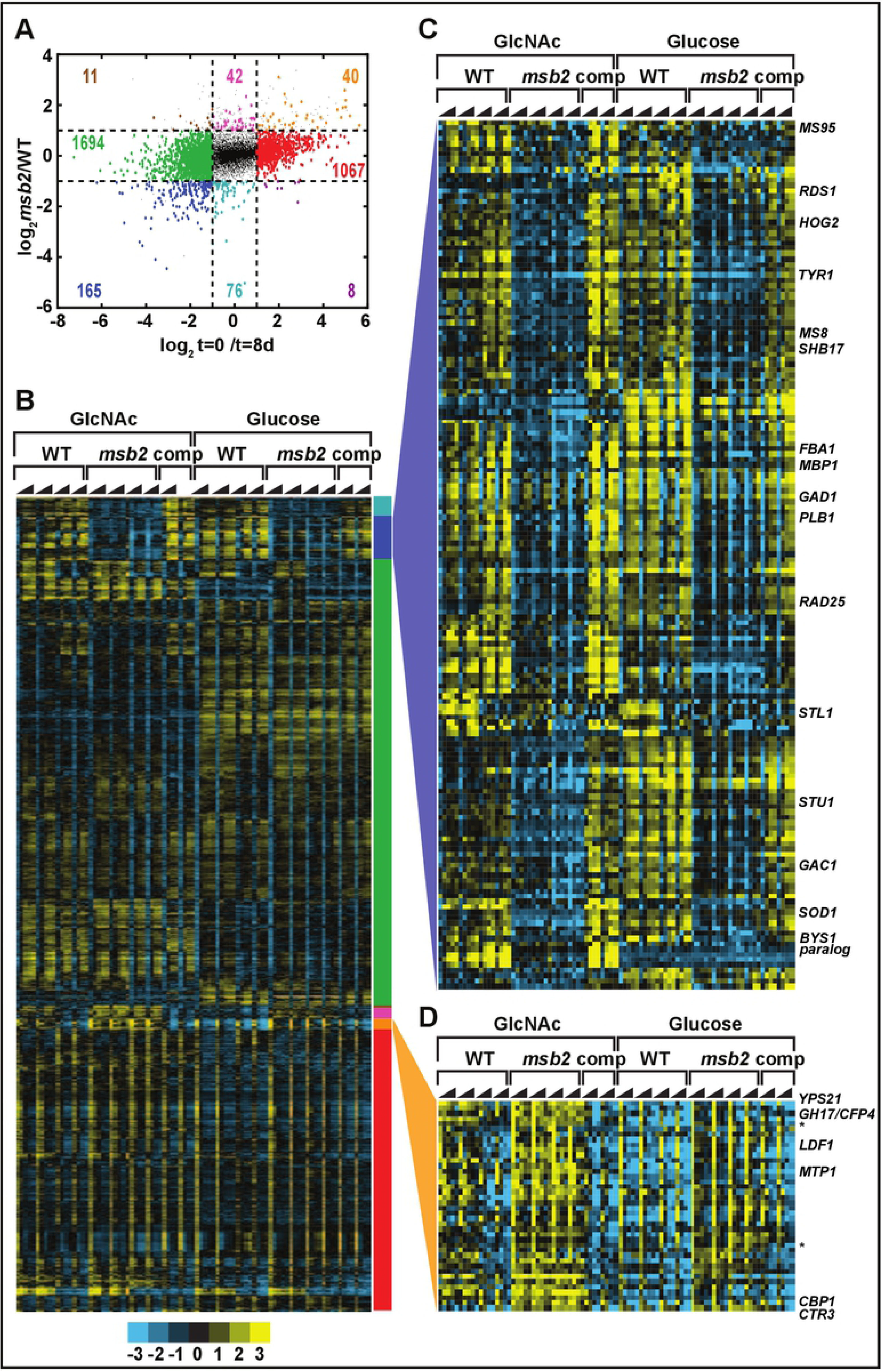
Classification of transcripts by dependence on *MSB2* and temperature. (A) Scatter plot of limma-fit parameters (as described in the Methods) for genotype (WT or *msb2* mutant) vs. time (day 0 or day 8). Transcripts significantly differential (at FDR = 5%) with at least a 2-fold change on one of these parameters are shown as larger, colored circles. Colored numbers indicate the number of transcripts in each class. (B) Heatmap of transcripts passing the significance and fold change criteria in A. Transcripts are grouped by the classification scheme of A, as indicated by the colored bar to the right of the heatmap. (C) Expanded view of the “filament-associated” class (165 genes) from 3A and B. (D) Expanded view of the “yeast-associated” class (40 genes) from 3A and B. Asterisks indicate putative knottin transcripts described in the text.

Analysis of the “filament-associated” gene set (Fig. 3C, S4 Data) revealed a number of orthologs of developmental regulators in other fungi. Among these was *FBA1*, which is a hyphal-specific gene that is upregulated in wild-type *H. capsulatum* at RT but not in *msb2*. It has an ortholog (*flbA*) in *Apergillus sp.* whose function has been elucidated as a regulator of conidiation [18]. While the cell morphologies of *Aspergillus spp* are distinct from those of *H. capsulatum* (for example, *Aspergillus spp* lack a yeast form) this result may suggest a central role for this gene as a regulator of morphological development in a wide range of fungi. Other potential regulators in the Msb2-dependent filament-associated gene set include the mitogen-activated protein kinase *HOG2* and the APSES transcription factor *STU1*; we assessed the role of these regulators in filamentation as described below.

Other genes in the filament-associated set have been flagged previously as showing hyphal-enriched expression in *H. capsulatum*. *MS8* and *MS95* have been shown to be highly hyphal-enriched in *H. capsulatum* in previous datasets through transcriptional profiling and ribosomal footprint analysis [4, 19, 20]. Here we observed that *MS8* and *MS95* are upregulated as early as 2 days at RT in a Msb2-dependent manner. Thus their expression is reliably correlated with growth at RT, but their molecular function and how it pertains to hyphal growth remains unclear. Notably, disruption of *MS8* has been reported to result in abnormally shaped hyphal cells [20]. Similarly, *TYR1*, which encodes a putative tyrosinase, has been observed as a highly abundant hyphal-enriched transcript in multiple studies [2–7]. Our data indicate that the transcriptional induction of *TYR1* after growth at room temperature is dependent on Msb2.

Of the 40 yeast-associated genes (Fig. 3D, S5 Data), 16 (40%) were previously identified as targets of the Ryp transcription factors, meaning that their promoters are associated with Ryp1, 2, 3, and/or 4. This is a significant enrichment over chance (p = 1.5e-7). Strikingly, all 16 are Ryp1-associated, with Ryp2, Ryp3, and Ryp4 associating with different subsets of these promoters. Since the Ryp program drives yeast-phase growth, and the *msb2* mutant is yeast-locked, these data suggest the hypothesis that the Ryp pathway is inappropriately active in the *msb2* mutant at room temperature. We previously observed that many known virulence factors are Ryp-associated [2]; consistent with this observation, the 16 Ryp-associated yeast-specific genes include 2 virulence factors, *CBP1* [21–24] and *CTR3* [25], as well as two genes required for efficient lysis of macrophages, *LDF1* and a gene previously designated as UA35-G3 [26]. Several others are putative secreted factors of unknown function, including the paralogs Yps21 [27] and GH17/Cfp4 [28, 29] as well as ucsf.hc_01.G217B.05476 which, along with UA35-G3, is predicted to be a secreted knottin (cystine knot proteins that show yeast-specific expression and may play a role in virulence in *H. capsulatum*) [4]. Notably, these yeast-associated genes, which are normally expressed by wild-type yeast cells at 37°C, are expressed in the yeast-locked *msb2* mutant even at room temperature, suggesting that their expression can be unlinked to temperature but remains linked to cell morphology.

### The MAP kinase *HOG2* is downstream of *MSB2* and is involved in the temperature-dependent regulation of filamentation

The function of Msb2 has been studied in a number of fungi [9–15, 30, 31]. These studies have supported the role of Msb2 as a transmembrane signaling molecule and more specifically, as one of the upstream signaling proteins in the *S. cerevisiae* high osmolarity glycerol (HOG) pathway [15, 32–34]. This pathway signals through the mitogen-activated protein kinase (MAPK) *HOG1* [35], which undergoes phosphorylation and translocation to the nucleus where it activates a transcriptional response to produce glycerol and restore osmotic balance. Given this precedent, we were interested in the possibility that Msb2 signals through a MAPK in *H. capsulatum*. Using phylogenetic analysis, we identified four MAPKs in *H. capsulatum* (Fig. 4A) and named them, in three cases, for their orthologs in other fungi (*HMK1, HOG1, SLT2*). The fourth MAPK is a paralog of Hog1 that we named Hog2. To determine if any of these were induced in hyphal cells, we examined the gene expression of these four MAPKs in our transcriptional profiling dataset. Whereas *SLT2, HMK1*, and *HOG1* did not show transcriptional induction in wild-type cells undergoing filamentation, *HOG2* was markedly induced in wild-type cells in an Msb2-dependent fashion (Fig. 4B). Additionally, *HOG2* was classified as a filament-associated transcript (Fig. 3C), and thus is a good candidate for a regulator of hyphal growth. To test whether *HOG2* is required for filamentation, we constructed a *HOG2* RNAi strain and assessed the morphology of these cells at RT. Isolates that showed efficient knockdown were yeast-locked, indicating that *HOG2* is required for filamentation at RT (Fig. 4C). Additionally, in the *MSB2* overexpression strain, which exhibits disruption of yeast morphology and filamentation at 37°C (Fig. 1E), we observed a significant increase in *HOG2* expression level compared to the control strain (Fig. 4D). Taken together, these data indicate that *HOG2* expression is increased in an Msb2-dependent manner and that *HOG2*, like *MSB2*, is necessary for filamentation in response to temperature.

**Fig. 4.**
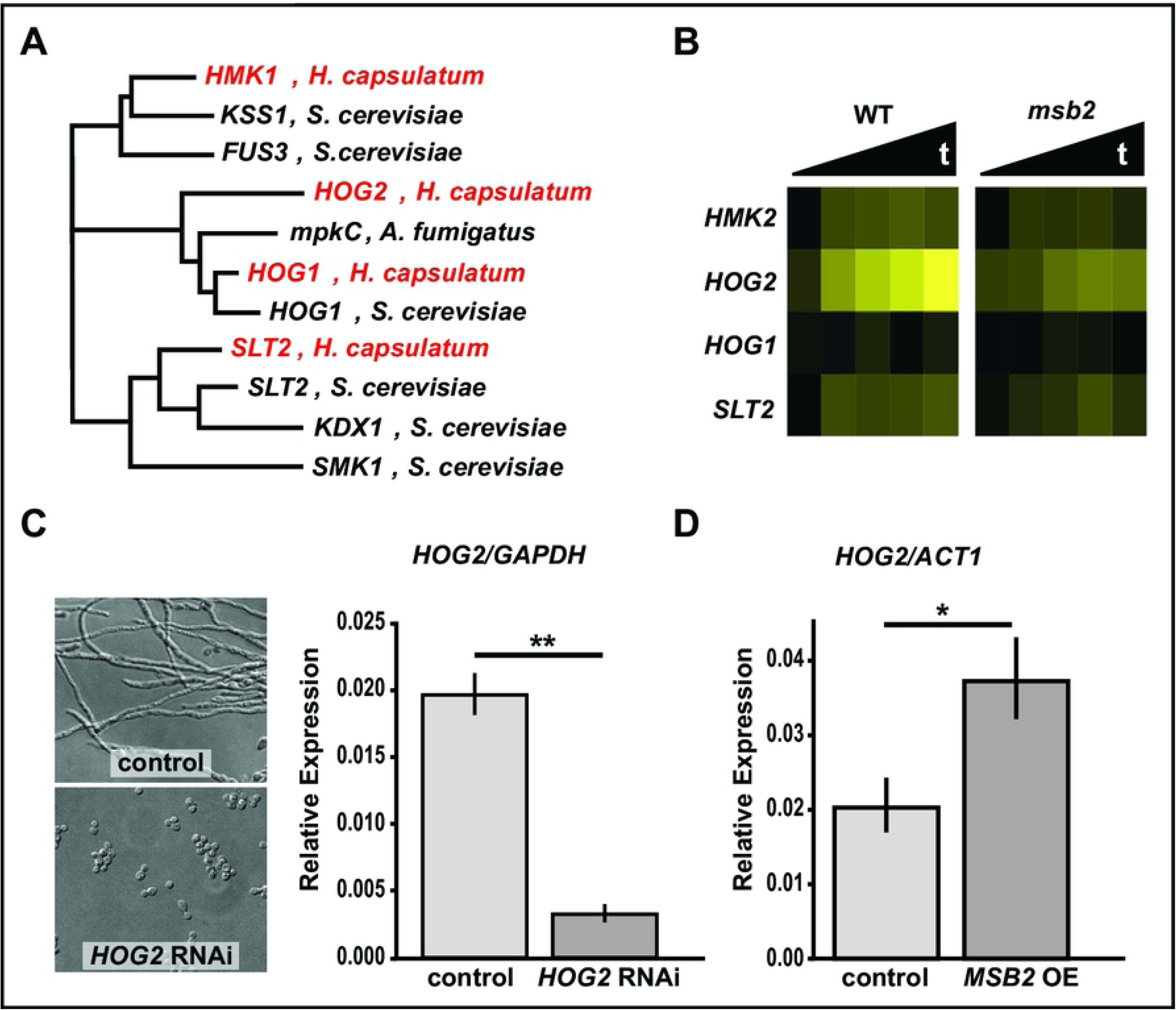
*HOG2* is a MAP kinase involved in filamentation in *H. capsulatum*. (A) FastTree maximum likelihood phylogeny of the 4 MAP kinases of *H. capsulatum* (shown in red) and their relation to other fungal MAP kinases. (B) Heatmap of *HMK2, HOG2, HOG1*, and *SLT2* data extracted from the RNAseq dataset. Values are log2 ratios from single GlcNAc cultures of WT or *msb2* at multiple time points after shift to RT, relative to a WT 37°C sample. (C-D) qRT-PCR data of *HOG2* transcript levels. (C) *HOG2* transcript levels in *HOG2* RNAi strain (P<0.01). Cell morphology of control and *HOG2* knockdown strains shown after 8 days at RT. (D) *HOG2* transcript levels in *MSB2* overexpression (OE) strain grown for 10 days at 37°C (P<0.05).

### The APSES transcription factor *STU1* is necessary and sufficient for filamentation

The Msb2-dependent filament-associated genes also include the transcription factor *STU1* (Fig. 3C). Stu1 orthologs have been thoroughly studied in species across the ascomycetes. The *Candida albicans* ortholog, *EFG1*, has long been known to promote filamentation, regulate other developmental transitions, and control gene expression [36–41]. StuA, the Stu1 ortholog in *Aspergillus spp.*, has been extensively studied as a developmental regulator [42, 43]. There are five APSES transcription factors in *H. capsulatum* [44], and Stu1 is the only one of the five that is transcriptionally induced as yeast-cells transition to filaments (Fig. 5A). As noted above, *msb2* mutant cells fail to induce *STU1* (Fig. 3C, Fig. 5A) but *STU1* expression is restored in the complemented strain at RT (Fig. 5B). To confirm *MSB2* is required for *STU1* expression in independent samples, *STU1* transcript level was measured by qRT-PCR in the control or *msb2* RNAi strain at RT and 37°C. We observed that *STU1* expression was significantly (p<0.001) depleted in the *msb2* knockdown strain at RT (Fig. 5C). To assess whether *STU1* is required for filamentation, we disrupted the *STU1* gene using CRISPR/Cas 9. We observed markedly reduced filamentation in the *stu1* mutant at RT (Fig. 5D), indicating that *STU1* is necessary for proper filamentation. Conversely, overexpression of *STU1* in wild-type *H. capsulatum* was sufficient to cause inappropriate filamentation at 37°C (Fig. 5E). Overexpression of *STU1* in the *msb2* mutant background was sufficient to restore the ability of the *msb2* mutant cells to form filaments at RT, and, like wild-type, inappropriate filamentation was observed at 37°C (Fig. 5E). Furthermore, cells harboring an *MSB2* overexpression plasmid not only displayed inappropriate filamentation at 37°C (Fig. 1E), but also showed inappropriate transcriptional induction of *STU1* at 37°C (Fig. 5F). Additionally, the *HOG2* RNAi strain, which is yeast-locked at RT (Fig. 4A), fails to induce *STU1* expression at RT (Fig. 5G). Taken together, our data strongly support the model that *MSB2* signals through *HOG2* and *STU1* to stimulate filamentation at RT (Fig. 5H).

**Fig. 5.**
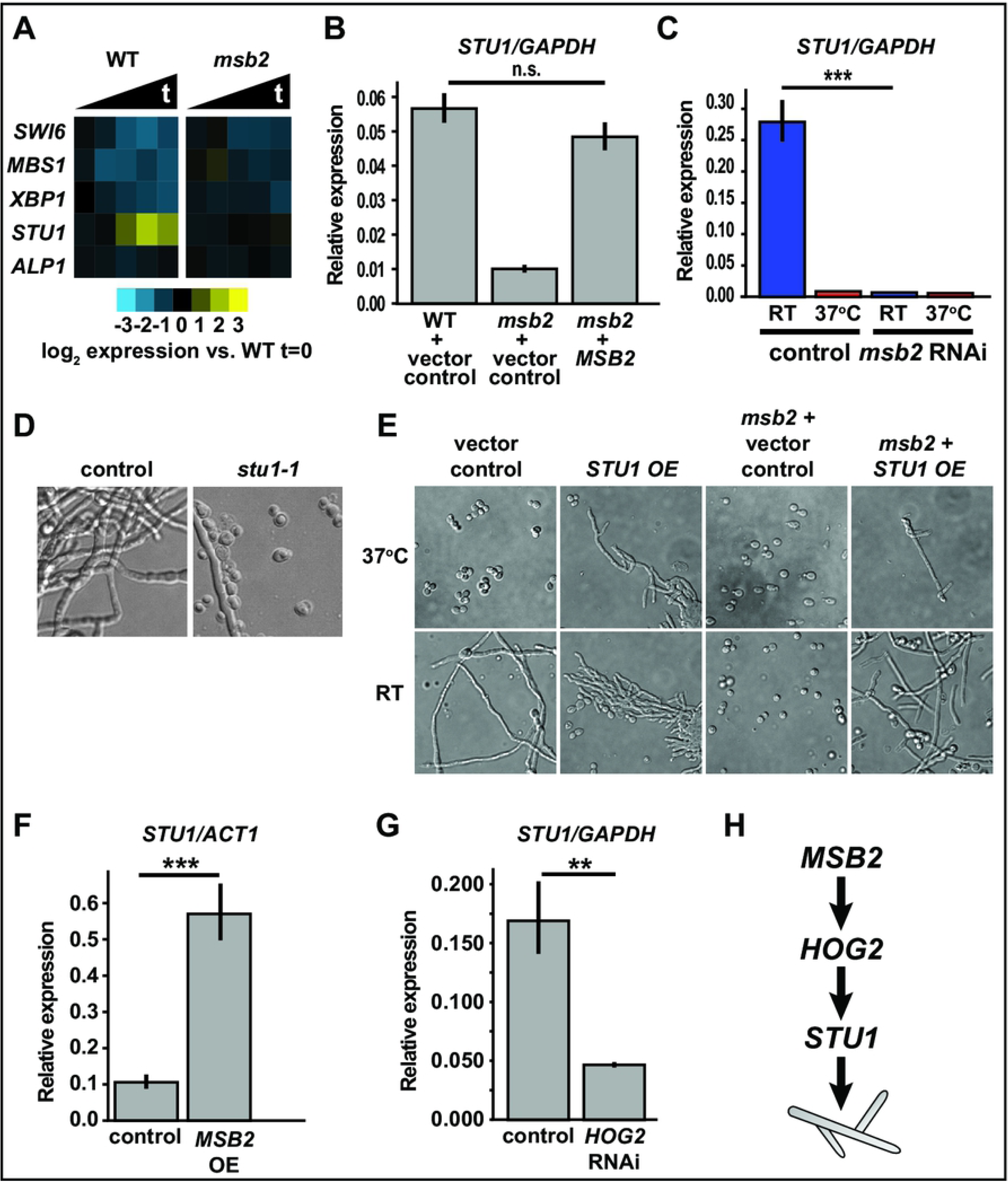
The conserved transcription factor *STU1* drives filamentation. (A) Heatmap showing transcription levels of APSES transcription factors in WT vs *msb2* extracted from the RNAseq dataset. Values are log_2_ ratios from single GlcNAc cultures of WT or *msb2* at multiple time points after shift to RT, relative to a WT 37°C sample. (B-C) qRT-PCR analysis of *STU1* transcript level (B) in WT compared to *MSB2* complemented strain after two weeks at RT (not significant); (C) in control compared to *msb2* RNAi strain after 2 weeks at RT (P<0.001). (D) Cell morphology of *stu1* disruption strain and WT after growth at RT for 8 days. (E) Cell morphology of strains produced using a control plasmid or *STU1* overexpression (OE) plasmid in wild-type or *msb2* background. Strains were grown at 37°C or RT. Control strains were imaged after 2 days 37°C or 5 days at RT and overexpression strains were imaged after 4 days at 37°C or RT. (F-G) qRT-PCR analysis of *STU1* transcript level in (F) *MSB2* overexpression strain (P<0.001); (G) *HOG2* RNAi strain (P<0.01). (H) Model showing the genetic relationship between *MSB2, HOG2*, and *STU1*.

### The “filament-associated” genes have conserved hyphal expression in ascomycete thermally dimorphic pathogens

As described above, our analysis defined filament-associated genes, which are expressed in an Msb2-dependent manner as wild-type cells undergo filamentation. Similarly, we defined yeast-associated genes, which are expressed in wild-type yeast cells at 37°C, and inappropriately maintain their expression in the *msb2* mutant at RT. As we demonstrated here, the filament-associated genes include regulators of filamentation such as *HOG2* and *STU1*, and the yeast-associated genes include previously identified virulence factors. We reasoned that if these morphological regulators and effectors have conserved function in other dimorphic fungi, then the “filament-associated” and “yeast-associated” regulons identified from our data should show conserved morphology-specific expression in these fungi.

In the case of filament-associated genes, this is indeed the case for three additional *Histoplasma* strains (G186AR, H88, and H143) and three additional thermally dimorphic ascomycetes (*Blastomyces dermatitidis, Penicillium marneffei*, and *Candida albicans*) (Fig. 6). In all cases where RNAseq data was available [3, 4, 45–47] (S6 Data), the distribution of yeast/hyphal expression for the orthologs of the filament-associated genes was biased towards showing hyphal expression rather than mimicking the distribution of the global transcriptome (p < .05, Wilcoxon test; S7 Data). In contrast, in *Ophiostoma novo-ulmi* (the causative agent of Dutch elm disease and the only other dimorphic ascomycete for which we were able to obtain RNAseq data [48]), the distribution of yeast/hyphal expression for the filament-associated regulon was indistinguishable from the rest of the transcriptome. It may be relevant that the dimorphic transition of *O. novo-ulmi* is both temperature-independent and rapid, with cells fully transitioning from yeast to hyphae within 27 hours.

**Fig. 6.**
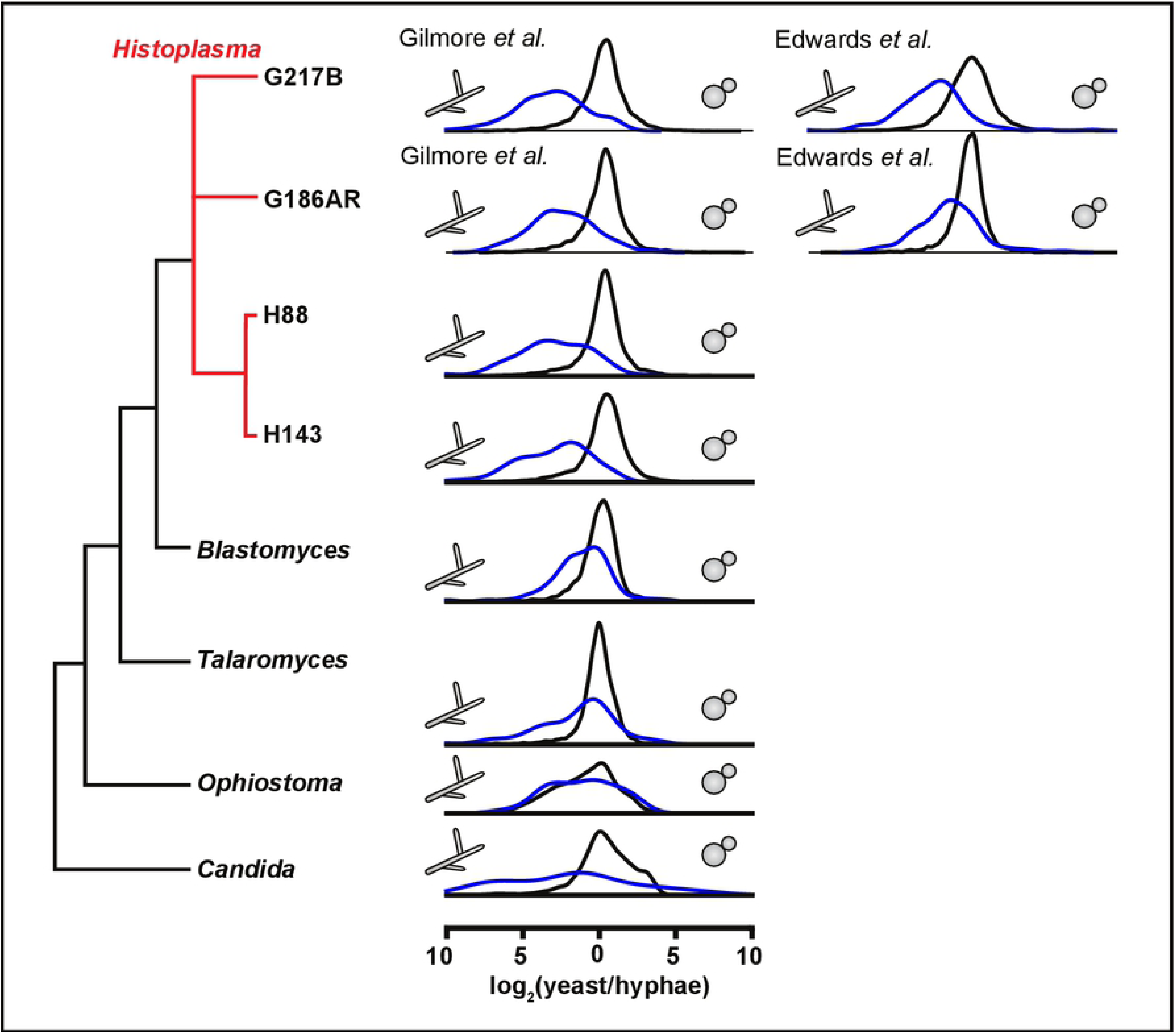
Msb2-dependent filament-associated genes are conserved across fungi. Distributions of log_2_(yeast/hyphal) expression ratios for orthologs of filament-associated genes from Fig. 3C (blue) or all other detected transcripts (black) for dimorphic ascomycetes with available RNAseq data. See Supp_comparative_sources.tdt data for data sources.

Notable genes with conserved hyphal expression in most or all of the thermally dimorphic fungi for which we could detect orthologs include *TYR1* (unique to the Ajellomycetacea), *MS95*, and *STU1* (S7 Data). We note that in *C. albicans, STU1* is homologous to the paralogs *EFH1* and *EFG1*. Whereas the *EFG1* transcript is not enriched in *C. albicans* hyphae relative to yeast cells in the RNAseq dataset we analyzed, the Efg1 protein has been shown previously to undergo post-translational regulation that affects its role in *C. albicans* hyphal morphology [49].

In contrast to the filament-associated genes, the yeast-associated genes had conserved morphologic expression only within *Histoplasma* using the same criteria as above. This may be due to *MSB2*-repressed yeast effectors that are unique to *Histoplasma* with respect to the dimorphic fungi listed above (e.g., *CBP1*).

### The Msb2 and Ryp programs oppose each other in response to temperature

Our data clearly show that *MSB2* and downstream genes such as *HOG2* and *STU1* are necessary and sufficient to drive a hyphal program. We previously identified four transcription factors, Ryp1-4, that are required for yeast-phase growth at 37°C, and show enhanced expression at 37°C over room temperature [2, 7, 8]. Since the *msb2* mutant is yeast-locked even at room temperature, we hypothesized that the *RYP* transcripts might be inappropriately expressed at RT in the *msb2* mutant, thereby allowing yeast-phase growth to persist. We compared *RYP* transcript levels by qRT-PCR in wild-type and *msb2* mutant strains at 37°C and RT (Fig. 7A). We determined that all four *RYP* transcripts are inappropriately maintained at higher levels in the *msb2* mutant at RT. This phenotype could be reversed by introducing a wild-type copy of *MSB2* into the mutant strain (Fig. 7B).

**Fig. 7.**
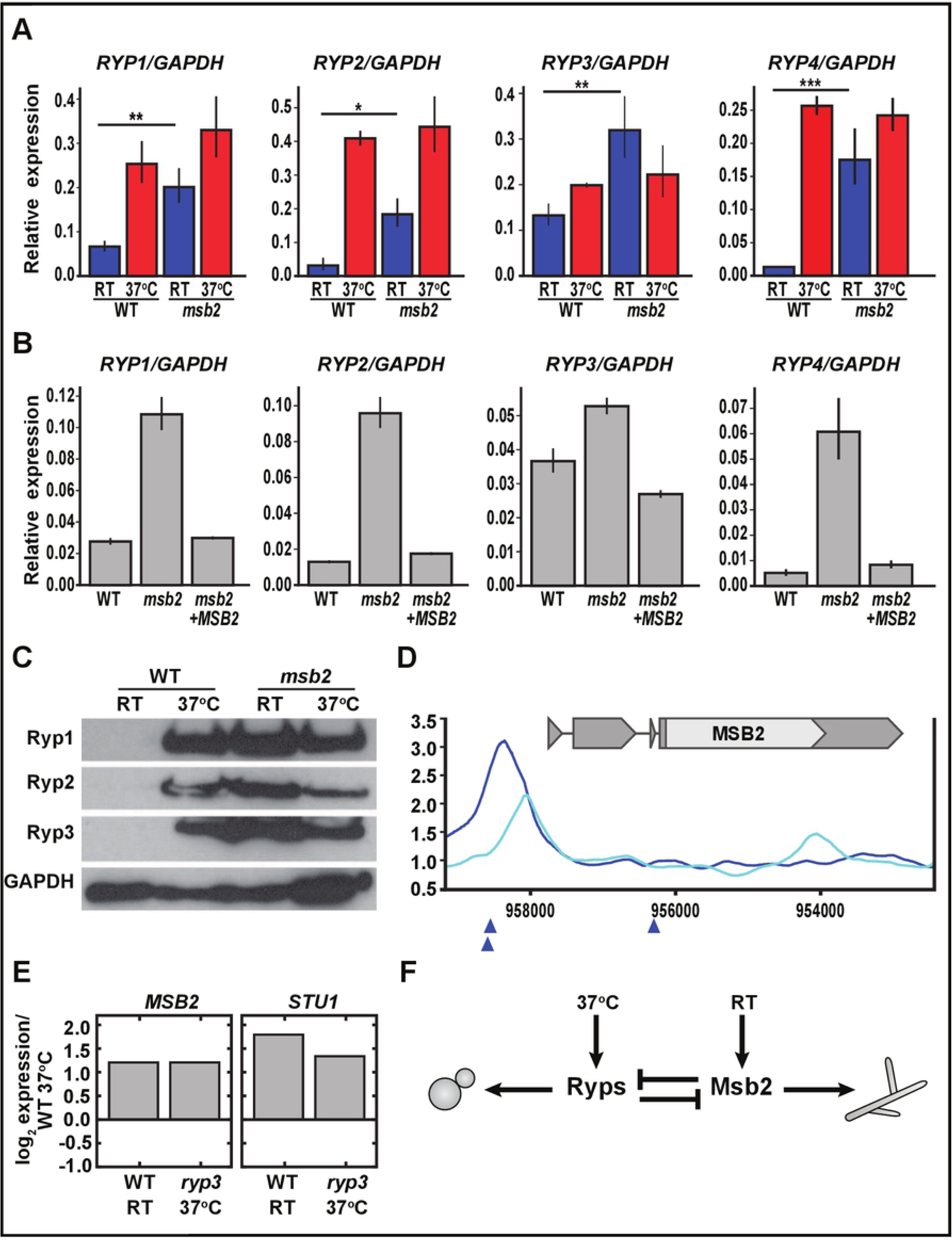
The Ryp and Msb2 programs antagonize each other in a temperature-dependent manner. (A-B) qRT-PCR analysis of *RYP1-4* transcript levels. (A) in WT strain compared to *msb2* after two weeks at RT: *RYP1* (P<0.01); *RYP2* (P<0.05); *RYP3* (P<0.01); *RYP4* (P<0.001); (B) in WT strain compared to *MSB2* complemented strain after two weeks at RT. (C) Western blots performed on protein extracted from cells grown either at 37°C or grown at RT for two weeks. GAPDH is a loading control. (D) Location of Ryp3 binding site upstream of *MSB2* transcript. Smoothed Ryp3 ChIP/input ratio ([2], GEO accession number GSE47341) is plotted for WT (dark blue) and *ryp3* control (light blue). Blue triangles indicate locations of Ryp2/Ryp3 associated Motif B [2]. (E) Microarray data [2] showing *MSB2* and *STU1* transcript levels in WT at RT and *ryp3* at 37°C, relative to corresponding transcript level in WT at 37°C. (F) Model showing the temperature-dependent relationship between the Ryps and Msb2.

As described above, we also noted that a number of the “yeast-associated” genes that are inappropriately expressed in the *msb2* mutant at RT are transcriptional targets of Ryp proteins. This observation implied that Ryp protein is inappropriately present at RT in the *msb2* mutant. We examined Ryp1, 2, and 3 proteins by Western blot and determined that these proteins are inappropriately expressed at RT in the *msb2* mutant (Fig. 7C). These data indicate that Msb2 is required to inhibit Ryp transcript and protein expression at RT.

We wondered if there was an analogous antagonistic relationship at 37°C, where Ryp proteins are required to inhibit *MSB2* transcript levels. We examined our previously published chromatin immunoprecipitation (ChIP-chip) data [2] and found a predicted Ryp3 binding site upstream of the *MSB2* open reading frame (Fig. 7D), suggesting that Ryp3 might repress *MSB2* transcript expression at 37°C. We examined our published microarray data for the *ryp3* knockdown strain [2]. In these experiments, as expected, the *MSB2* transcript showed upregulation at RT relative to 37°C in wild-type cells. Notably, the *MSB2* transcript levels at 37°C in the absence of *RYP3* were at levels similar to the wild-type strain at RT (Fig. 7E), indicating that *RYP3* is required to prevent transcriptional induction of *MSB2* at 37°C. Moreover, *RYP3* is also required to inhibit *STU1* expression at 37°C (Fig. 7E), indicating that the Ryp program inhibits expression of multiple members of the Msb2 hyphal program at 37°C. Taken together, we conclude that the Ryp program and the Msb2 program oppose each other in a temperature-dependent manner such that the Ryp program dominates at 37°C whereas the Msb2 program dominates at RT, ultimately allowing regulation of cell shape by temperature (Fig. 7F).

## Discussion

Thermally dimorphic fungal pathogens display an exquisite response to temperature that governs their ability to thrive in either the environment of the soil or the mammalian body. At environmental temperatures, these fungi grow in a multicellular hyphal form that is suited to survival in the soil. In response to mammalian body temperature, these organisms switch to a host form (unicellular yeast cells in the case of *Histoplasma*) that expresses virulence factors that are important for disease progression. Although the existence of this response is thought to be critical to the pathogenesis of endemic fungi such as *Histoplasma, Blastomyces, Paracoccidioides*, and *Coccidioides* spp.[50], the molecular basis of this dichotomous response to temperature is not understood. Here we uncover a pathway required for the filamentation response to environmental temperature, and show that this pathway acts in a mutually antagonistic manner with the regulators that promote the morphologic response to mammalian body temperature. Thus regulation of thermal dimorphism represents a binary choice, where the fungus transitions between two different states in response to temperature by means of opposing regulatory pathways.

A number of cell fate decisions in both microbes and multicellular organisms are controlled by bistable switches, where feedback mechanisms drive the system into one of two possible dynamic steady states. In the case of *Histoplasma* thermal dimorphism, we showed previously that the Ryp proteins form an interlocking circuit that promotes yeast-phase growth [2]. All four Ryp proteins bind upstream of Ryp1, 2, and 4, which is thought to create a positive feedback loop that contributes to the marked increase of Ryp protein levels at 37°C. Additionally, ribosomal profiling data suggest that the translational efficiency of Ryp1 and Ryp2 is higher at 37°C than at room temperature [4]. Notably, Ryp3 associates with the upstream region of the *MSB2* gene at 37°C, which is associated with decreased accumulation of the *MSB2* transcript [2] (Fig. 7). Thus several mechanisms cooperate to enhance Ryp protein levels at 37°C. In contrast, at room temperature, the abundance of Ryp proteins is markedly decreased. During room temperature growth, *MSB2* is required for the morphologic switch from yeast-phase cells to hyphal cells, and molecularly, *MSB2* is required for the reduction in Ryp transcript and protein levels (Fig. 7). Notably, overexpression of *MSB2* is sufficient to trigger morphologic changes even at 37°C (Fig. 1), suggesting that levels of *MSB2* are a critical determinant of cell shape. Taken together, our data show that in wild-type cells, inhibition of the Ryp pathway by an Msb2-dependent mechanism is likely to be a key step in the switch from yeast to filaments in response to temperature. Notably, in *Candida spp*, an antagonistic regulatory relationship between Wor1 (an ortholog of the *H. capsulatum* Ryp1 protein) and Efg1 (an ortholog of the *H. capsulatum* Stu1 protein) is critical for controlling the switch between distinct developmental states [36, 51–53].

*In vitro* experiments show that morphologic transitions in the thermally dimorphic fungi occur over the timescale of days rather than hours, suggesting that sustained changes in temperature are required to shift cell shape. It is attractive to speculate that *H. capsulatum* is integrating a temperature signal over time to enable specific responses to the homeostatic temperature environment of the host while ignoring transient spikes in environmental temperature. In molecular terms, the precise temporal parameters that govern a switch between dominance of the Ryp pathway and dominance of the Msb2 pathway are unknown. Further experimentation may reveal that sustained but not transient temperature shifts are required for one antagonistic program to fully disrupt and reset the other. Ultimately, multiple molecular mechanisms could contribute to the role of temperature in these cell-fate decisions.

This study is the first to elucidate components of the pathway that transduce the morphogenetic response to environmental temperature in *H. capsulatum*. The role of Msb2, a transmembrane signaling mucin, in temperature response is particularly intriguing. Msb2 has long been known to regulate the filamentous growth and osmosensing pathways in *Saccharomyces cerevisiae* [11, 14] as well as to influence morphogenesis in *C. albicans* [10] and appressorium development in *Ustilago maydis* [30]. Additionally, recent work showed that Msb2 regulates responses to temperature stress in *C. albicans* [54]. It has become clear that proteolytic cleavage of the extracellular domain of Msb2 is a critical step in signaling [12, 55]. It is attractive to speculate that this type of cleavage could be triggered by temperature shift and contribute to Msb2-dependent responses in *H. capsulatum*.

A number of studies have shown that the transcriptional profile of 37°C-grown yeast cells and RT-grown hyphae are quite distinct [2–7], although the function of the majority of differentially expressed transcripts is unknown. One of the most informative aspects of the *MSB2* analysis is that the compact Msb2 regulon distinguishes genes that are key to filamentation or virulence from the high background of differentially expressed factors. As shown in this work, the *msb2* mutant appears morphologically as yeast at room temperature, and we initially predicted that its transcriptional profile would resemble that of wild-type yeast cells. In actuality, during RT growth, the *msb2* mutant transcriptome resembles that of wild-type hyphae despite its yeast-phase morphology. Specifically, the majority of the ∼1870 genes that are differentially expressed by wild-type hyphae at RT are also expressed by *msb2* mutant yeast at RT (Fig. 3A), and the majority of the ∼1100 genes that are differentially expressed by wild-type yeast cells at 37°C fail to maintain their expression at RT in the yeast-form *msb2* mutant (Fig. 3A). The fact that the *msb2* mutant grows as yeast at RT but looks transcriptionally like hyphae is in contrast to the *ryp* mutants, which appear both morphologically and transcriptionally as hyphae at 37°C [2, 7]. Interestingly, there is a set of yeast-phase genes whose expression is decreased after shift to room temperature in wild-type cells, but inappropriately maintained in the *msb2* mutant. This set of 40 genes is enriched for direct transcriptional targets of the Ryp transcription factors, suggesting that the Ryp proteins, which are present at inappropriately high levels at RT in the *msb2* mutant, are able to activate expression of a subset of their targets. Importantly, our experiments reveal that expression of these genes appears to be tightly coupled to growth in the yeast form rather than growth at 37°C. It seems highly significant that this gene set contains known virulence factors and small, secreted proteins that may be well positioned to manipulate events within host cells. Thus the transcriptional analysis of the *msb2* mutant defined 40 yeast-associated transcripts that are of high interest for potential roles in virulence.

In an analogous fashion, the *msb2* mutant also defined a core set of 167 filament-associated transcripts that fail to be expressed in the yeast-locked mutant at RT when compared to wild-type cells, suggesting that only a fraction of the RT transcriptional program is required for the developmental process of filamentation. This result is analogous to findings in *C. albicans*, where the majority of the transcriptional response to filamentation conditions is dispensable for hyphal formation [47, 56]. We reasoned that the *H. capsulatum MSB2* regulon was likely to contain regulators and effectors of the hyphal growth program, and indeed confirmed that *HOG2* and *STU1* are required for filamentation. Notably, a recent publication examining APSES transcription factors in *H. capsulatum* showed that knockdown of *STU1* resulted in a defect in aerial hyphae production [44], which agrees with our data indicating that *STU1* is involved in hyphal growth. A number of other putative regulatory factors are members of the *MSB2* regulon, and their functions remain to be analyzed. When this Msb2-dependent filamentous gene set was compared to other *Histoplasma* strains and more distantly related fungi, we found that, on average, this gene set displayed conserved, filament-enriched expression. This suggests that the *MSB2* regulon is conserved across a diverse group of fungi and that the genes involved in filamentation are shared in multiple fungal species. Notably, this filament-associated gene set may highlight factors that have important regulatory or effector roles in the hyphal growth program of other fungi. We are particularly interested in ucsf_hc.01_1.G217B.09523, which lacks an ortholog in *C. albicans*, but was present with conserved hyphal enrichment in all other fungi in our dataset, including *O. novo-ulmi*. This gene is orthologous to fungal *Metarhizium anisopliae* Crp2 and plant *Arabidopsis thaliana* atRZ-1a, sharing their N-terminal RRM_1 RNA binding domain and glycine rich C-terminus. Crp2 and atRZ-1a are thought to provide cold resistance by acting as RNA chaperones, and are sufficient to confer this resistance when heterologously expressed in *Saccharomyces cerevisiae* [57] *and an Escherichia coli* cold-sensitive mutant [58], respectively. Intriguingly, two other RRM_1 family members with broad RNA binding activity have distinct effects on hyphal growth in the basidiomycete dimorphic fungus *Ustilago maydis*: Rrm4 mediates proper polar growth filaments via long-distance transport of mRNAs along hyphae while Grp1 may act as an RNA chaperone as well as an accessory component in endosomal mRNA transport [59]. The intriguing relationship between expression of putative RNA-binding factors and thermal dimorphism has yet to be fully explored, but may shed light on how adaptation of fungi to different environments is linked to morphologic changes.

Finally, the identification of regulators of the environmental differentiation program has provided insight into critical antagonists of the host program. Elucidating the molecular switches that control the host program may lead to novel tactics to target virulence strategies in thermally dimorphic fungi.

## Materials and Methods

### *H. capsulatum* Strains

*Histoplasma capsulatum* G217B *ura5-* (WU15; referred to in this work as wild-type) was used for the insertional mutagenesis screen, which was performed using the *Agrobacterium tumefaciens* strain LBA1100 containing the plasmid pRH5b as described [7]. All strain manipulations were done in the *H. capsulatum* G217B *ura5-* background. Strains used in this paper can be found in Table S3.

### Media

Wild-type G217B, complementation, and RNAi strains were grown in liquid or solid *Histoplasma* Macrophage Medium (HMM) [60]. G217B *ura5-* and insertional mutants were grown in HMM supplemented with 2.5mg/ml uracil. Broth cultures were grown at 37°C with 5% CO2 on an orbital shaker at 120 –150 rpm. For yeast-form colonies, plates were grown in a humidified incubator at 37°C with 5% CO2. Plates grown at room temperature were wrapped individually in Parafilm and placed in a 25°C standing incubator in a Biosafety Level 3 facility. Cells were thawed fresh from frozen stock and passaged on HMM agarose plates every 1–2 weeks for up to 2 months. 100mM N-acetylglucosamine (Sigma-Aldrich) was used as the primary carbon source in HMM where indicated.

### Screen for yeast-locked mutants

A small pilot screen was performed to identify yeast-locked mutants. *H. capsulatum* G217B *ura5-* yeast were co-cultured with *Agrobacterium* carrying plasmid pRH5b, which has a T-DNA element with a hygromycin marker, allowing for selection of insertion mutants as previously described [7]. The transformants were plated at a dilution aiming for 50 colonies per plate and were incubated at 37°C, 5% CO2 until colonies appeared. After 25 days, the plates were transferred to a room temperature incubator in the Biosafety Level 3 laboratory. 92 colonies were observed for colony morphology. After 13 days, 2 of the 92 colonies retained yeast morphology while the others had filamentous projections.

### Mapping the T-DNA insertion in the *msb2* mutant

The location of the T-DNA insertion was determined from whole-genome sequencing. The SG1 strain (the *msb2* mutant) and the control strain were grown up from frozen glycerol stocks onto HMM agarose plates supplemented with uracil. Liquid cultures were inoculated for each strain and passaged twice at 1:25 dilution in HMM + uracil. Two days after the second passage, the cells were collected via centrifugation. Genomic DNA of the strains was isolated using the Gentra Puregene Yeast Kit (Qiagen). Libraries were made using the Nextera XT DNA Library Prep Kit (Illumina).

The libraries were multiplexed with 7 other *Histoplasma* genomic DNA libraries and sequenced on an Illumina HiSeq 4000, yielding 21,038,128 paired-end 101-mer reads for ∼106x coverage of the ∼40MB G217B genome. Bowtie2 was used to map these read pairs to a joint index of the T-DNA plasmid sequence (pRH5b) plus the 11/30/2004 version of the G217B genome assembly from the Genome Sequencing Center at Washington University (GSC) as mirrored at http://histo.ucsf.edu/downloads/. The location of the T-DNA insert was determined by filtering for discordantly mapped read pairs with one mate in the T-DNA and one mate in the reference genome. Discordant pairs consistently mapped to the T-DNA left border and HISTO_ZL.Contig1131 or the T-DNA right border and HISTO_DA.Contig93, indicating a single insertion event resulting in a genomic rearrangement. A crude model of the HISTO_ZL.Contig1131/T-DNA/HISTO_DA.Contig93 hybrid sequence was constructed based on the discordantly mapped reads, erring on the side of including extra sequence from HISTO_ZL.Contig1131 and HISTO_DA.Contig93. Bowtie2 was then used to remap the reads to the model sequence, allowing the exact junction coordinates to be determined from the alignment of reads spanning the genomic/T-DNA boundaries. This gave a final model of the hybrid sequence (Fig. 1B, bottom), which was confirmed by Bowtie mapping of RNAseq reads from the SG1 mutant to the hybrid sequence (Fig. 1B, green coverage lines).

It is evident from this RNAseq mapping that *MSB2* is transcribed in the SG1 mutant, with transcription initiating from a site in the T-DNA. The resultant extended 5’ region of the transcript introduces a new open reading frame (pink box in Fig. 1B), which we hypothesize acts as an upstream ORF that inhibits Msb2 translation.

### Construction of plasmids

Sub-cloning was performed in *E. coli* DH5α. All knockdown constructs were generated by amplifying a 500 bp region to target a particular gene for RNA interference as previously described [2]. The *MSB2* gene was targeted by amplifying G217B genomic DNA using OAS5486-87 (primers are described in Table S2). The fragment was cloned into the pDONR/Zeo vector using Gateway Cloning Technology (Invitrogen), generating the plasmid pLR04. The final episomal RNAi vector was produced by recombining pSB23 [2] with pLR04, generating plasmid pLR11. The *HOG2* RNAi strains were generated with the same methodology. The 500 bp region used to target *HOG2* was amplified using OAS5005-06 and the PCR product was cloned into pDONR/Zeo, generating plasmid pSB420. This plasmid was then recombined with pSB23 to produce pSB430.

The overexpression construct for *STU1* was produced by placing the *ACT1* promoter upstream of the *STU1* coding sequence. This construct was generated by amplifying the *ACT1* promoter using OAS2594 and OAS2587, *STU1* using OAS2586 and OAS2592, and the *CATB* terminator using OAS2591 and OAS2593. The fragments were joined using the overlap extension PCR cloning method [61]. The construct was recombined with pDONR/Zeo using Gateway Cloning Technology (Invitrogen) to produce the plasmid pTM1. pTM1 was recombined with the destination plasmid pDG33 to produce the final plasmid pTM2. The overexpression construct for *MSB2* was produced by placing the GAPDH promoter upstream of the *MSB2* coding sequence. OAS5476-77 was used to amplify the coding sequence by PCR. Using Gateway Cloning Technology (Invitrogen), the PCR fragment was cloned into the pDONR/Zeo plasmid to produce pLR09. This plasmid was recombined with pSB234, which is an episomal plasmid carrying the GAPDH promoter upstream of the Gateway cassette, making it compatible with the Gateway cloning system (Invitrogen). pLR02 and pSB234 were recombined to produce pLR09.

The *MSB2* complementation construct was generated by using the primers OAS5711-12 to amplify the gene and its native promoter. This fragment was cloned into pDONR/Zeo using Gateway Cloning Technology (Invitrogen) to produce the plasmid pLR01. pLR01 was recombined with pDG33 to produce pLR09.

The plasmids used in this study are described in Table S1 and primers are described in Table S2. To generate RNAi, complementation, or overexpression strains, episomal plasmids were linearized, electroporated into the relevant yeast strain, and plated onto HMM agarose plates without uracil to select for *URA5*-containing transformants. The strains used in this study are described in Table S3.

### Construction of the *stu1* mutant strain

We made an episomal version (pBJ219) of an *Agrobacterium*-based Cas9 targeting plasmid [62] (kind gift of Bruce Klein). pBJ219 is described in a separate manuscript (Joehnk and Sil, in preparation). In brief, a guide RNA targeting the first exon of *STU1* was introduced into pBJ219, which carries a fungal codon-optimized Cas9 along with ribozyme sequences required for the excision of the RNA sequence [62]. The plasmid was introduced into the G217B *ura5-* strain by electroporation and transformants were selected on HMM plates lacking uracil. Single colonies were isolated from individual transformants, genomic DNA was prepared, and the genomic sequence at the *STU1* locus was analyzed using TIDE, an online tool measuring the frequency of indels in a mixed population [63]. After three sequential passages, we colony-purified an isolate with an insertion of an A at position 23 relative to the ATG, resulting in an early stop codon after amino acid 13.

### Imaging

Imaging was performed on a Zeiss AxioCam MRM microscope using DIC at 100X magnification.

### Culture conditions for expression profiling

Samples were prepared in two distinct experiments and sequenced in three distinct sequencing runs. In the first experiment, two biological replicates each of *Histoplasma capsulatum* G217B *ura5* (WU15) or an *msb2* mutant derived from WU15 (SG1) were grown in liquid culture with glucose or GlcNAc as the carbon source at 37°C. After taking t = 0 samples, the cultures were moved to RT and additional samples were taken at 2, 6, 8, or 10 days. One sample was not usable for sequencing, giving a total of 2*2*2*5-1 = 39 samples. The GlcNAc samples were sequenced in part of a lane of one HiSeq 4000 run (batch 1) and the glucose samples were sequenced in part of a lane of a second run (batch 2). Because these samples were grown at the same time, they were treated as a single batch in our limma analysis. In the second experiment, two biological replicates each of WU15, SG1, or SG1 complemented with the wild-type *MSB2* gene (SG1cMSB2) (resulting from transformation with pLR08) were grown in liquid culture with glucose or GlcNAc as the carbon source at 37C. In this second experiment, WU15 and SG1 were transformed with the empty vector control (pLH211) so that the parental, mutant, and complemented strains were comparable. After taking t = 0 samples, the cultures were moved to RT and additional samples were taken at 2, 6, or 8 days, for a total of 2*3*2*4 = 48 samples. These samples were sequenced in a full lane of a HiSeq 4000 run (batch 3).

### Fungal cell collection

*H. capsulatum* cultures were collected by filtration with a disposable filtration apparatus either in the Biosafety Level 2* laboratory for cultures grown at 37°C or in the Biosafety Level 3 laboratory for cultures grown at RT. At each time point, 10 mL of fungal culture was passed through a disposable filter apparatus (Thermo Fisher), the cells were scraped off with a cell scraper, placed into a conical tube with 1 mL Trizol reagent (Qiazol) and flash frozen in liquid nitrogen. Samples were stored at −80°C until all time points were collected.

### RNA extraction

Total RNA was extracted from fungal cells using a Trizol-based RNA extraction protocol. Frozen, resuspended pellets of cells were incubated at RT for 5 minutes to thaw. The lysate was subjected to bead beating (Mini-Beadbeater, Biospec Products) followed by a chloroform extraction. The aqueous phase was then transferred to an Epoch RNA column where the filter was washed with 3 M NaOAc and then with 10 mM TrisCl in 80% EtOH. DNase (Purelink) treatment was used to remove any residual DNA and the filters were washed again with NaOAc and TrisCl before eluting the RNA in nuclease-free water. RNA quality was determined with a High Sensitivity DNA Bioanalyzer chip (Agilent).

### mRNA isolation

20 µg of total RNA was purified for mRNA through polyA selection with Oligo-dT Dynabeads (Thermo Fisher) as described in the manufacturer’s protocol. Ribosomal RNA depletion was confirmed with an RNA 6000 Nano Bioanalyzer chip (Agilent Technologies). Samples were acceptable for library preparation at 0-4% rRNA.

### RNAseq library preparation

Libraries for RNAseq were prepared using the NEB Next Ultra Directional RNA Library Prep Kit (New England Biolabs). Individual libraries were uniquely barcoded with NEBNext Multiplex Oligos for Illumina sequencing platform (New England Biolabs). Average fragment size and presence of excess adapter was determined with High Sensitivity DNA bioanalyzer chip from Agilent Technologies. Libraries had an average fragment length of 300-500 bp. The concentration of the individual libraries was quantified through Qubit dsDNA High Sensitivity Assay (Invitrogen). 5 ng of each library was pooled into three final libraries (batches in S1 Data) and run on a High Sensitivity DNA Bioanalyzer chip to determine the average fragment size of the final pooled samples. The final libraries were submitted to the UCSF Center for Advanced Technology for sequencing on a Illumina HiSeq4000 sequencer.

### RNAseq data analysis

For quality control, data exploration, and to quantify yields (mapping statistics in S1 Data), samples were aligned with Bowtie version 1.1.2 [64] to the 11/30/2004 version of the G217B genome assembly from the Genome Sequencing Center at Washington University as mirrored at http://histo.ucsf.edu/downloads/, invoked as:

zcat SAMPLE.fastq.gz | bowtie -tS -p 4 -k 1 HcG217B -\

| samtools view -bS -o SAMPLE.bam

Transcript abundances were quantified based on version ucsf_hc.01_1.G217B of the *H. capsulatum* G217B transcriptome (S5 Data of [4],

http://histo.ucsf.edu/downloads/ucsf_hc.01_1.G217B.transcripts.fasta)

Relative abundances (reported as TPM values [65]) and estimated counts (est_counts) of each transcript in each sample were estimated by alignment free comparison of k-mers between the preprocessed reads and assembled transcripts using KALLISTO version 0.43.0 [66], invoked as:

kallisto quant -i ucsf_hc.01_1.G217B.idx -t 4 -b 100 \

--single --rf-stranded -l 250 -s 50 \

-o SAMPLE.kallisto SAMPLE.fastq.gz

Further analysis was restricted to transcripts with TPM >= 10 in at least one sample.

Differentially expressed genes were identified by linear regression on four independent factors using LIMMA version 3.26.1 [67, 68].

Specifically, KALLISTO est_counts were rounded to the nearest integer and imported as counts per million (CPM) in a DGEList in R. Samples were TMM normalized with calcNormFactors and VOOM [69] was used to estimate the mean/variance trend.

Samples were classified on four factors (S1 Data): batch (1 or 3), genotype (wild-type or *msb2*), media (GlcNAc or glucose), and time (0, 2d, 6d, or 8d). For each transcript, the observed counts were fit to a model assuming a mean transcript abundance plus independent corrections for each factor using lmFit. Shrinkage of the individual gene variances over the full data was applied with eBayes, and significantly differential genes were identified by extracting genes with a significantly non-zero genotype and/or t = 8d coefficient (at 5% FDR) with an effect size of at least 2x (absolute log2 fold change >= 1) using topTable.

The core R invocations were:

dge <-DGEList(counts)

dge <-calcNormFactors(dge)

time <-as.factor(time)

genotype <-as.factor(genotype)

media <-as.factor(media)

batch <-as.factor(batch)

d <-model.matrix(∼time+genotype+media+batch) v <-voom(dge, d)

fit <-lmFit(v, d)

fit <-eBayes(fit)

topTable(fit, coef=“genotypeMSB2”, n = 20000, lfc=1, p.value = .05) topTable(fit, coef=“time8”, n = 20000, lfc=1, p.value = .05)

### Comparative Transcriptome Analysis

For all ascomycete yeast/hyphal dimorphs with available RNAseq data (S6 Data), reads were downloaded from the SRA. Corresponding transcriptome sequences were downloaded from the sources indicated in S6 Data and indexed for kallisto. Transcripts were quantified for each sample with kallisto, invoked as:

kallisto quant -i TRANSCRIPTOME.idx -t 4 EXTRA RUNS

where TRANSCRIPTOME.idx is the appropriate index file, RUNS are gzipped FASTQ files for all runs corresponding to the given sample, and EXTRA are additional flags for appropriate experiment-specific handling of single-ended and strand-specific reads, as given in the “extra flags” column of S6 Data. Further analysis was restricted to transcripts with TPM >= 10 in at least one sample. Log2(TPM) values were averaged across biological replicates for each morphology and then subtracted to give log2(yeast/hyphae) differential expression values (S7 Data). *Histoplasma* ortholog groups were taken from S13 Data from [4]. Othologous genes between G217B and the remaining species were determined by InParanoid version 1.35 [70].

We tested the null hypothesis that log2(yeast/hyphae) values for orthologs of the G217B filament-associated genes were drawn from the same distribution as the remaining genes with G217B orthologs using a two sided Wilcoxon rank sum test. Kernel density estimates of these distributions are plotted in Fig. 6 using R’s density function.

### Expression level by qRT-PCR

Relative gene expression was observed using qRT-PCR. RNA extraction was performed as described above and cDNA was prepared by priming 2 µg DNase-treated total RNA with oligodT/pdN9 and dNTPs. Samples were shifted to RT and treated with 1 µl RNaseOUT Recombinant Ribonuclease Inhibitor (ThermoFisher) and 0.5 µl Thermo Scientific Maxima H Minus Reverse Transcriptase. Negative controls included samples that were processed without the addition of reverse transcriptase. Samples with reverse transcriptase were diluted 1:50 and samples without reverse transcriptase were diluted 1:10. qPCR reactions were set up using 96-well plates with FastStart Universal SYBR Green Master (Roche) and 1.5 µM primers (found in Table S2), and plates were read using a Mx3000P machine (Stratagene) and analyzed using MxPro software (Stratagene).

### Protein extraction

Organic fractions from Trizol RNA extractions were stored at −20°C until protein extraction was performed. To remove DNA from the organic fraction, 100% ethanol was added and samples were centrifuged to pellet DNA. The phenol-ethanol supernatant containing the proteins was transferred to a fresh tube containing isopropanol and centrifuged at 4°C to pellet the protein precipitate. The pellet was washed with 0.3M guanidine thiocyanate in 95% ethanol and centrifuged at 4°C. This wash step was repeated for a total of 3 times. 100% ethanol was added to the protein, which was then vortexed and incubated for an additional 20 minutes at RT. The protein was pelleted by centrifugation and air-dried at RT. Finally, the pellet was resuspended in urea lysis buffer (9M Urea, 25mM Tris-HCl, 1mM EDTA, 1% SDS, 0.7M β-mercaptoethanol). If necessary, the pellet was incubated at 50°C for 10-20 minutes to successfully resuspend it in the urea buffer. Once resuspended, the sample was boiled, centrifuged, and the supernantant, containing the protein was transferred to a new tube for quantification by the Pierce 660 nm assay with added ionic detergent compatibility reagent (ThermoFisher).

### Expression level by Western blot

Western blotting was performed with polyclonal peptide antibodies against either Ryp1, Ryp2, or Ryp3; antibodies were described previously [2]. Following quantification of protein, 10-30 µg was resuspended in a total of 20 µL of urea lysis buffer and 6 µL NuPAGE LDS Sample Buffer (Novex). The samples were boiled and electrophoresed on a 10-well NuPAGE 4-12% BT SDS-PAGE gel (Novex) in MOPS running buffer at 150V. The protein was then transferred to a nitrocellulose membrane at ∼40 V for 2 hours. The membrane was incubated with blocking solution (1 g milk powder in 100 mL wash buffer (0.1% Tween-20 in PBS)) for an hour and then incubated in the primary antibody in wash buffer overnight at 4°C. Primary antibody dilutions were: α-Ryp1 (1:10,000), α-Ryp2 (1:2,500), α-Ryp3 (1:5,000), α-GAPDH (1:1,000). The blot was washed and secondary antibody (for Ryp proteins: Goat α-rabbit HRP (GenScript) 1:1,000, for GAPDH: Goat α-mouse HRP (ThermoFisher) 1:1,000) was added to the blot for 1 hour at RT followed by another wash. Protein bands were detected using chemiluminescence according to the manufacturer’s instructions (SuperSignal West Pico kit (ThermoFisher)).

### MAP Kinase Phylogeny

MAPK protein sequences were obtained from SGD (https://www.yeastgenome.org/; HOG1=YLR113W, SMK1=YPR054W, KDX1=YKL161C, STL2=YHR030C, KSS1=YGR040W, FUS3=YBL016W), AspGD (http://www.aspgd.org/; mpkC=AN4668), and HistoBase (http://histo.ucsf.edu/downloads/; HOG1=ucsf_hc.01_1.G217B.05737, HOG2=ucsf_hc.01_1.G217B.11562, HMK1=ucsf_hc.01_1.G217B.10227, SLT2=ucsf_hc.01_1.G217B.02764) and aligned with PROBCONS 1.12 [71]. A phylogeny was generated from the PROBCONS alignment with FastTree 2.1.7 [72].

### Statistical Analysis

qRT-PCR data analysis was performed as described [73] with significant differences determined by the t-test using propagated standard errors. Growth curves were graphed with Prism (GraphPad Software).

### Other software and libraries

We wrote custom scripts and generated plots in R 3.2.3 [74] and Python 2.7.12, using Numpy 1.11.0 [75] and Matplotlib 1.5.1 [76]. Jupyter notebooks [77] and JavaTreeView [78] were used for interactive data exploration.

## Acknowledgements

We are grateful to Chad Rappleye and Bruce S. Klein for generously sharing reagents and information. We thank Bevin English for helpful advice and suggestions, and Alexander Johnson and Joseph DeRisi for their guidance. Eric Chow and the UCSF Center for Advanced Technology provided invaluable advice on library preparation and sequencing. We also thank Davina Hocking Murray for help with figure design and preparation, as well as Hiten Madhani, Suzanne Noble, and members of the Sil and Noble labs for helpful discussions.

## Supporting Figure Legends

**Fig. S1.**
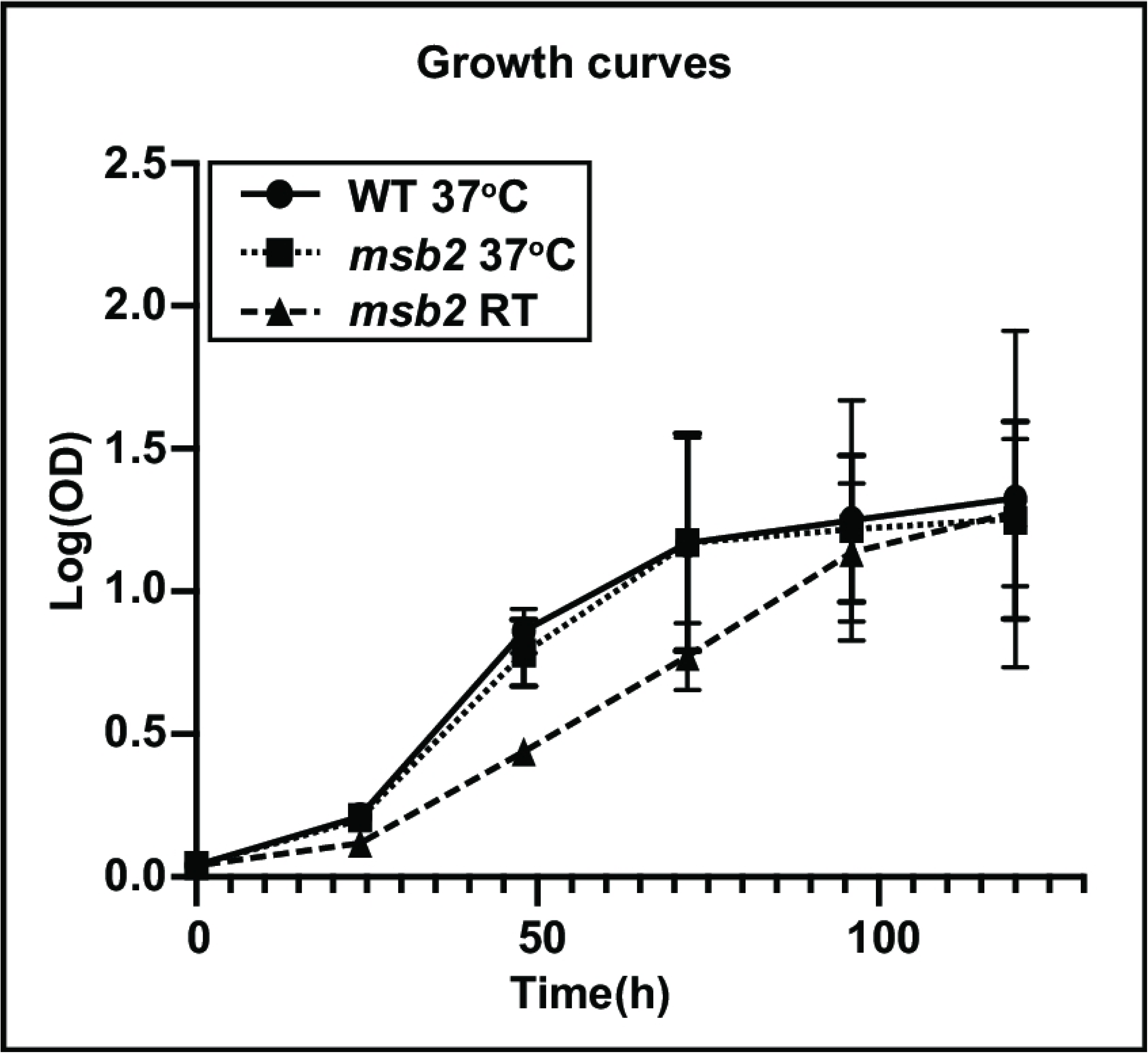
*msb2* yeast grown at RT continue to divide. A growth curve of WT and mutant yeast at 37°C is compared to mutant yeast grown at RT.

**Fig. S2.**
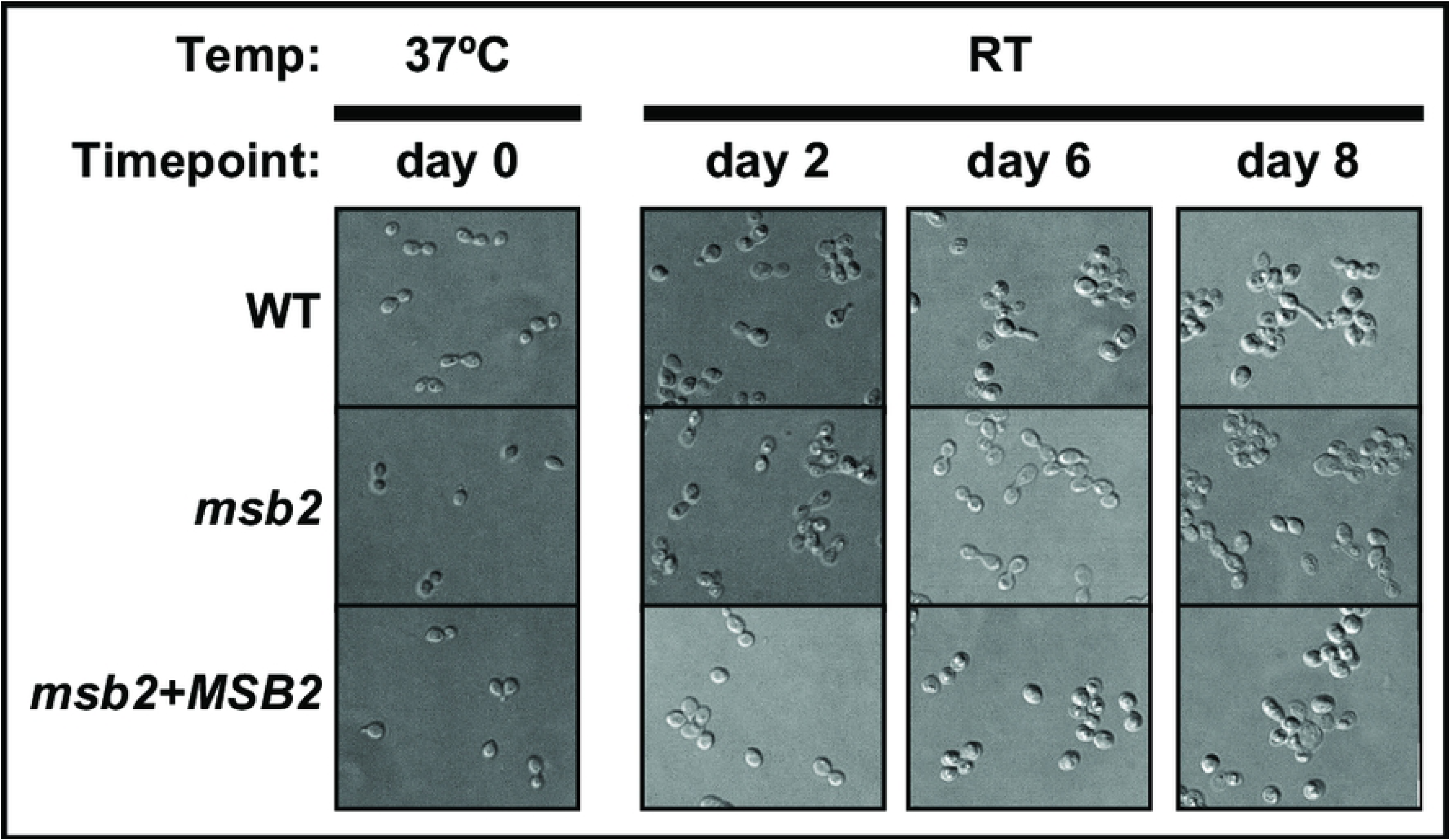
Cells grown for RNAseq timecourse in glucose as carbon source do not transition to hyphae by 8 days after the shift to RT. Cells were grown at 37°C (d0) or RT (d2-8) and imaged to observe morphology.

## Supporting Data Legends

All supporting data is formatted as Excel-compatible tab-delimited text. Expression profile data sets (S2, S3, S4, S5, and S7) are additionally compatible with the JavaTreeView extended CDT file format.

**S1 Data. Sequenced samples.** Columns are unique sample name; the four parameters used in limma fitting, *viz*.: genotype (G217B = wild type, SG1 = *msb2* mutant, SG1cMSB2 = complemented *msb2* mutant), time (days after transfer to RT), media (GlcNAc or glucose), and batch (where batches 1 and 2 were treated as a single batch while fitting); as well as total read depth and number of reads mapped to the G217B assembly by Bowtie.

**S2 Data. Kallisto estimated counts and limma fit values for time course expression profiles.** Each row is a transcript, with the UNIQID column giving the ucsf_hc.01_1.G217B systematic gene name. The NAME column gives short names taken from Data S13 of [4] with additional names and corrections based on human curation. The next 87 columns give KALLISTO estimated counts for each transcript in each sample (with column headings corresponding to sample names in Data S1). Additional annotation columns are G217B predicted gene name, *H. capsulatum* conserved enrichment, and signal peptide, all taken from Data S13 of [4]; and ChIP-chip based association with the Ryp transcription factors, taken from Table S2 of [2]. The final 7 columns give the limma fit parameters, in units of log2(CPM), and the corresponding limma probabilities that the non-intercept parameters differ from 0 are given as p(parameter).

**S3 Data. Genes significantly differential in response to temperature and/or *msb2* mutation.** Each row is a transcript, with UNIQID and NAME, and annotation columns as in S2 Data with an additional “class” column indicating the eight expression classes from Fig 3A and 3B. The remaining columns give log2(CPM) values for each sample (with column heading corresponding to sample names in Data S1). These values are TMM normalized and each row is mean centered, such that positive and negative values correspond to samples with expression above or below the average expression for a given transcript. As in Fig 3B, transcripts are grouped by class and sorted by the limma fit MSB2 dependence within each class.

**S4 Data. Yeast associated genes.** Subset of S3 Data corresponding to genes significantly up in both WT/*msb2* and t = 0/t = 8d.

**S5 Data. Filament associated genes.** Subset of S3 Data corresponding to genes significantly down in both WT/*msb2* and t = 0/t = 8d.

**S6 Data. Data sources for comparative transcriptome analysis.** Each row corresponds to a profile plotted in Fig. 6. Columns give a shorthand name, taxonomic details (genus, species, strain), data accession (SRA) and reference (PMID), sequencing layout (paired or single), extra flags supplied to KALLISTO (see Methods), the specific SRA sample accessions used for the yeast or hyphae expression profiles, and the source of the transcriptome FASTA file used to build the KALLISTO index.

**S7 Data. Comparative transcriptome profiles.** Each row corresponds to an *H. capsulatum* G217B transcript with UNIQID, NAME, G217B predicted gene, and class as in Data S3. Ortholog columns give the corresponding transcript names for other species (empty for cases with no mapped ortholog). Y/H columns give KALLISTO estimated log2(yeast/hyphae) expression ratios corresponding to sample names in Data S6.

